# H3BERTa: A CDR-H3 specific language model for antibody repertoire analysis

**DOI:** 10.1101/2025.11.03.686198

**Authors:** Chiara Rodella, Thomas Lemmin

**Affiliations:** Institute of Biochemistry and Molecular Medicine, University of Bern, Bühlstrasse 28, 3012 Bern, Switzerland; Graduate School for Cellular and Biomedical Sciences (GCB), University of Bern, Mittelstrasse 43, 3012 Bern, Switzerland

**Keywords:** CDR-H3, antibody, protein language model, machine learning, broadly neutralizing antibodies (bnAbs), repertoire analysis

## Abstract

Antibodies are central to immune defense and therapeutic design, yet predicting which sequences confer functional activity remains challenging. Deep learning models trained on full variable regions often struggle due to sparse experimental data, signal dilution from conserved framework residues, and the extreme diversity of hypervariable loops. The heavy-chain complementarity-determining region 3 (CDR-H3) is the most variable segment shaping antigen specificity and driving immune diversity. Here, we present H3BERTa, a transformer-based language model trained solely on CDR-H3 sequences, to test whether this short region alone encodes enough biologically meaningful information. H3BERTa embeddings recapitulate biologically relevant sequence features, including J-gene usage and inferred B-cell maturation state. We further show that pseudo-perplexity profiles can be used to analyze repertoires, distinguishing healthy from HIV-1–derived sequences and suggesting measurable immune response signatures. Finally, these embeddings can support classifiers for broadly neutralizing antibodies (bnAbs) using limited labeled sequences, demonstrating their potential for accelerating antibody discovery. Together, our results indicate that the CDR-H3 region alone encodes a rich immunological signature, which H3BERTa robustly captures, providing a focused computational tool for analyzing repertoire diversity and informing antibody engineering.

## 1 Introduction

The identification of antigen-specific antibodies within the vast and diverse B-Cell Receptor (BCR) repertoire remains a major challenge in immunology, vaccinology, and therapeutic antibody discovery. Advances in Next-Generation Sequencing (NGS) technologies have enabled the high-throughput profiling of immune repertoires, generating datasets that comprise millions of antibody sequences from individual donors [1–3]. However, the vast majority of these sequences do not exhibit specificity to a given target antigen, and only a small fraction possess high-affinity or broadly reactive properties [4, 5]. Among these rare antibodies, broadly neutralizing antibodies (bnAbs) are of particular interest due to their ability to recognize conserved epitopes across diverse viral strains [6, 7]. bnAbs have been shown to neutralize challenging pathogens such as HIV-1 [8, 9], influenza [7, 10], and coronaviruses[6], and have emerged as promising candidates for passive immunization, vaccine design, and therapeutic intervention. Unfortunately, their low prevalence in natural repertoires and the difficulty of linking sequence to function continue to limit their discovery. Furthermore, time consuming computational approaches are required in order to efficiently identify these rare but biologically significant molecules from large-scale sequencing data.

Among the most studied examples of bnAbs are those targeting Human Immunodeficiency Virus type 1 (HIV-1) [11]. These antibodies exhibit the extraordinary ability to neutralize a broad spectrum of HIV-1 strains by binding to conserved regions of the viral envelope glycoprotein, such as the CD4-binding site, the V1/V2 apex, and the membrane-proximal external region [8, 9]. Beyond their direct therapeutic and prophylactic potential, HIV-1 bnAbs have provided critical insights into B cell maturation, immune tolerance, and the co-evolution of host and virus [12, 13]. However, these antibodies are typically generated in only a small subset of infected individuals [14], often after prolonged periods of antigenic stimulation and extensive somatic hypermutation. As a result, their identification through experimental methods remains difficult, time-consuming, and resource-intensive [15].

Despite these obstacles, recent advances in machine learning and structural modeling have begun to reshape how we study antibody repertoires. Classical approaches, such as clonal lineage tracing, sequence motif detection, or alignment-based similarity metrics, offer valuable insights but often lack scalability and generalizability to new antigens or hosts [16, 17]. Currently, language models trained on large corpora of antibody sequences have shown promise in learning context-aware representations that encode structural, functional, and evolutionary features without requiring explicit supervision [2, 18–20]. These models, inspired by architectures from natural language processing, enable powerful sequence embeddings that have been applied to antigen specificity prediction, paratope modeling, and immune repertoire classification.

However, many of these models are trained on full-length variable regions, requiring extensive data for downstream tasks, more computational resources and potentially obscuring functionally relevant signals due to conserved framework residues. In this study, we propose a more targeted strategy centered on the immunoglobulin heavychain Complementarity-Determining Region 3 (CDR-H3), a short but highly variable segment that plays a central role in antigen recognition. CDR-H3 is known to contribute disproportionately to binding specificity and diversity, and has been shown to encode signatures of immune selection, clonal expansion, and somatic mutation [1]. We hypothesized that focusing exclusively on CDR-H3 would be sufficient to extract immunologically meaningful representations, especially in settings where annotated data is sparse or full-length sequences are unavailable. To test this hypothesis, we developed H3BERTa, a transformer-based language model trained specifically on the CDR-H3 region. Despite its narrow input scope, H3BERTa learns biologically relevant representations that capture features such as gene usage and B cell maturation state. Leveraging these representations, we show that the model can be used to distinguish repertoire-level differences between healthy individuals and HIV-infected patients and explored the potential for a classifier that uses CDR-H3 embeddings to identify candidate HIV-1 broadly neutralizing antibodies. This approach enables repertoire-wide screening using only short sequence fragments, and demonstrates strong potential for accelerating antibody discovery, particularly in data-limited settings.

## 2 Results

The immunoglobulin (Ig) heavy-chain CDR-H3 is critical for antigen specificity and commonly used in repertoire analysis, often through methods relying on simple features like length or sequence similarity [21, 22]. However, such approaches offer a limited view of the complex biological information encoded within this highly variable region. We hypothesized that, despite its short length, the CDR-H3 region encodes sufficient immunological information to support nuanced repertoire analysis using advanced deep learning techniques.

To investigate this, we developed H3BERTa, a transformer-based language model leveraging the RoBERTa architecture [23] and comprising approximately 86 million trainable parameters. H3BERTa was pretrained on approximately 18 million CDRH3 sequences obtained from the Observed Antibody Space (OAS) database [24]. This extensive dataset was specifically curated to include IgG and IgA isotypes from healthy, unvaccinated donors, thus providing a broad and biologically meaningful foundation reflecting major functional antibody classes in systemic and mucosal immunity (Figure 1A). IgG antibodies are typically the most abundant in serum and mediate a wide range of effector functions, whereas IgA antibodies play a crucial role at mucosal surfaces [25].

**Fig. 1.**
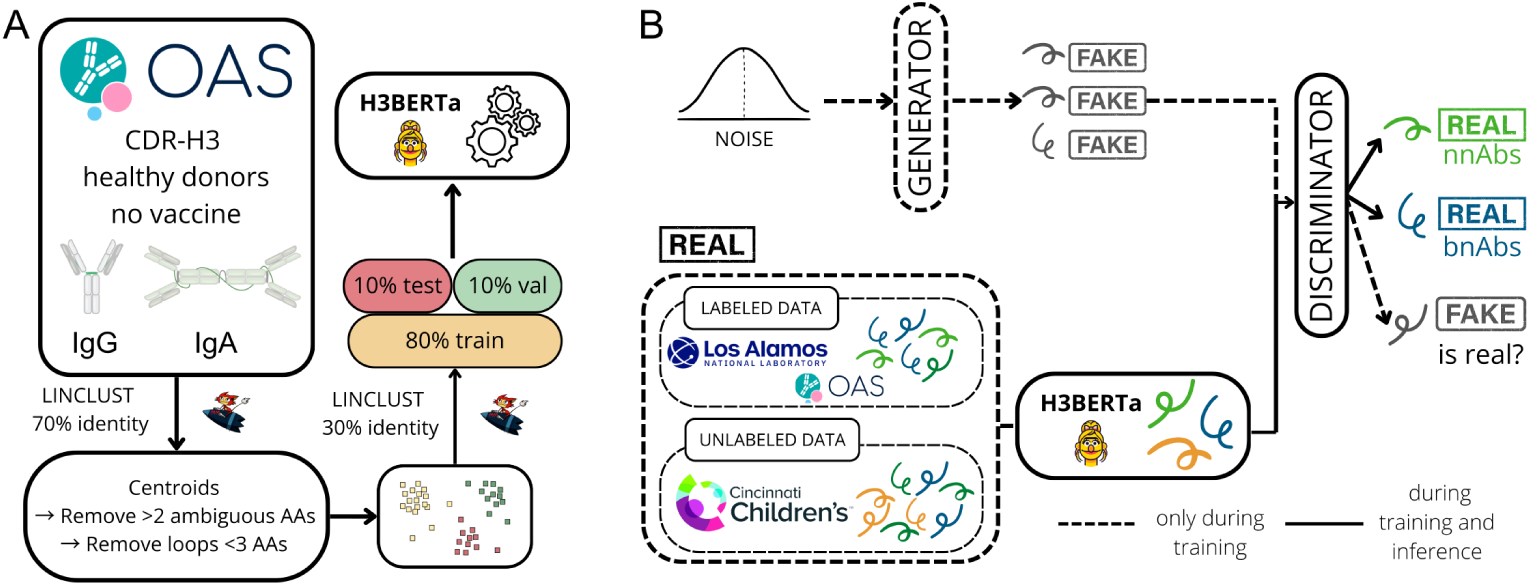
Overview of the H3BERTa deep learning framework and its application to broadly neutralzing antibodies (bnAbs) classification. (A) H3BERTa pre-training. Data preparation and training pipeline for H3BERTa using diverse CDR-H3 sequences from the Observed Antibody Space (OAS) database. (B) Semi-supervised bnAb classification with GANH3BERTa. The discriminator is jointly trained to distinguish real from synthetic embeddings and to classify sequences as broadly neutralizing (bnAb) or non-neutralizing (nnAb) antibodies. Posttraining, the discriminator is used to predict bnAbs in antibody repertoires.

H3BERTa was trained for 113 epochs, converging with a final masked language modeling loss of 1.58. To further assess the biological relevance of its amino acid predictions, we calculated average pairwise substitution scores using the BLOSUM62 matrix[26], with an average score of 2.55. While not tailored to CDR-H3 diversity, this metric provides a general baseline and suggests that the model captures biologically plausible substitutions rather than relying solely on exact matches.

To assess whether the CDR-H3 region alone captures sufficient immunological signal for repertoire-level analysis, we compared H3BERTa to AntiBERTa [27], a transformer-based model pre-trained on full variable regions of immunoglobulin heavy chains. This comparison allowed us to evaluate how a minimal, CDR-H3 centric representation performs relative to models trained on more extensive sequence input. A UMAP visualization of H3BERTa embeddings revealed a repertoire organization primarily influenced by the J-gene segment (Figure 2A). This demonstrates H3BERTa’s capacity to organize sequences based on intrinsic genomic signals associated with the CDR-H3 recombination process. In contrast, embeddings from AntiBERTa showed a stronger clustering by V-gene segment (Figure 2B), reflecting the model’s access to the framework and other CDR regions, where V-gene identity is a dominant signal.

**Fig. 2.**
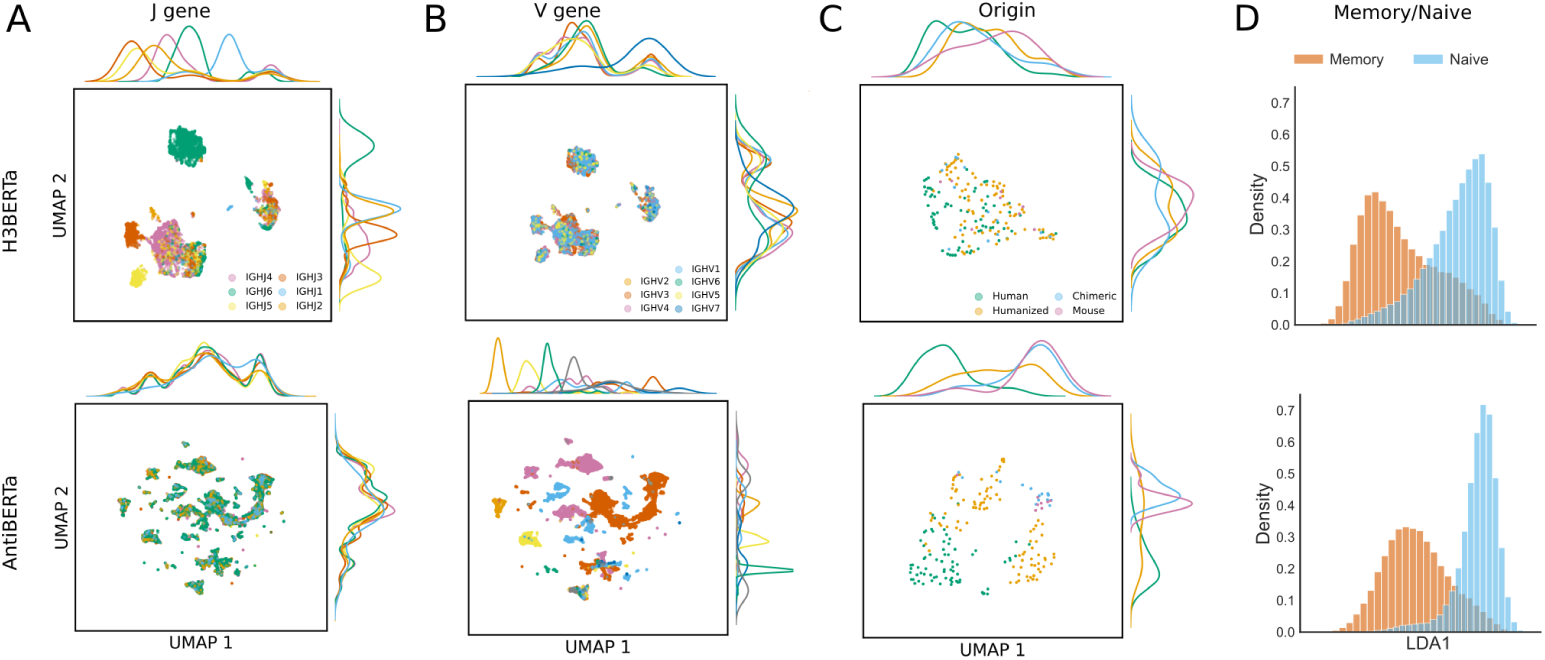
Comparison of antibody sequence embeddings from H3BERTa and AntiBERTa models. Sequence embeddings generated by H3BERTa (top row) and AntiBERTa (bottom row), visualized using UMAP or Linear Discriminant Analysis (LDA). For UMAP panels, marginal distributions along each axis are shown as kernel density ridges. Each point represents a unique antibody heavy-chain sequence. UMAP projection of embeddings, colored by (A) IGHJ gene segment and (B) IGHV gene segment. (C) UMAP projection of embeddings from therapeutic antibodies, colored by annotated species origin: Human (green), Humanized (orange), Chimeric (light blue), and Mouse (pink). (D) LDA projection of embeddings from naive (blue) and memory (orange) B cell sequences.

Next we evaluated each model’s ability to capture species-specific features in therapeutic antibody sequences by generating embeddings for antibodies from the Thera-SAbDab database [28], which includes approved and clinical-stage antibodies annotated by species of origin. AntiBERTa embeddings exhibited clear clustering by species (Figure 2C), reflecting the presence of species-distinctive sequence features in the variable domain. These include framework residues that are less frequently mutated during affinity maturation and thus preserve phylogenetic and germline-derived differences between species and antibody formats. Since AntiBERTa processes the entire variable region, it captures these invariant residues and species-specific motifs that distinguish engineered from native antibodies. In contrast, H3BERTa embeddings showed a more diffuse distribution, with only partial separation for some human antibodies. This could suggest that weak species-related signals are still partially encoded within the CDR-H3 region.

In addition to gene-specific signals, we sought to determine whether H3BERTa’s embeddings could capture the B cell maturation state, a critical aspect of repertoire evolution. Given that UMAP projections were predominantly influenced by gene segment distributions, we employed Linear Discriminant Analysis (LDA) to specifically delineate maturation-associated features. By applying LDA to a labeled dataset of naive and memory B cell sequences, both H3BERTa and AntiBERTa consistently generated highly separable, bimodal distributions (Figure 2D), with only marginal overlap between the naive and memory peaks. This strong separation indicates that despite different input scopes, both models effectively capture distinct sequence features characteristic of B cell maturation, further validating the biological richness encoded within their learned representations.

Finally, we computed the Peusdo-Perplexity (PPL) of both models on a healthy donor BCR repertoire [29], which estimate the likelihood of a sequence under the model. PPL reflects how well a sequence conforms to the learned statistical patterns of the model’s training data and thus serves as a proxy for repertoire typicality. The PPL scores were moderately correlated between H3BERTa and AntiBERTa (Pearson *r* = 0.41, Spearman *ρ* = 0.60; Figure S1), indicating that despite differences in training input, both models capture partially overlapping aspects of sequence composition.

### 2.0.1 Characterizing bnAb repertoires using H3BERTa

Having established that H3BERTa can organize antibody repertoires in an unsupervised manner based on intrinsic sequence features, we next investigated whether it could also detect functional differences between biologically distinct repertoires. HIV broadly neutralizing antibody (bnAb) repertoires were selected as a test case, since these arise from chronic antigen exposure and prolonged affinity maturation, processes expected to imprint distinct sequence characteristics on the CDR-H3 region [30, 31]. If H3BERTa’s embeddings capture these subtle immunogenetic signals, then they should allow discriminating between bnAb and healthy donor repertoires.

CDR-H3 sequences were analyzed from a published dataset [29], which includes 4.4 million sequences from 42 healthy donors and 4.2 million sequences from 46 bnAb donors. For our analysis, one representative donor from each group was selected. In the H3BERTa embedding space, both donors showed a comparable overall organization, with UMAP projections consistently revealing three distinct clusters corresponding to IGHJ6, IGHJ4, and IGHJ3 usage (Figure S2), consistent with the germline-driven structure observed in earlier analyses. No clear clustering pattern was observed when stratifying by V gene usage. However, no overt group-level separation was visible, suggesting that bnAb-associated differences, if present, were probably more subtle than gene-segment–driven patterns and would require more sensitive statistical comparisons.

To probe these differences, we turned to the distribution of pseudo-perplexity values assigned by H3BERTa to sequences within each repertoire. For each donor, the full PPL distribution was computed across all CDR-H3 sequences and quantified interindividual differences using pairwise Wasserstein distances. This metric, also known as Earth Mover’s Distance, is sensitive to shifts in both mean and distributional shape, making it well-suited for quantifying repertoire-level dissimilarities.

Although the overall distributions of PPL appeared visually similar between healthy and bnAb repertoires (Figure S3), hierarchical clustering based on pairwise Wasserstein distances revealed a broad separation between the two groups (Figure 3A). This suggests that subtle but systematic shifts in the shape or spread of individual repertoire PPL profiles are sufficient to distinguish cohort-level differences. Consistent with this, the average Wasserstein distance between healthy and bnAb repertoires was higher than the average within-group distances (Figure S3A).

**Fig. 3.**
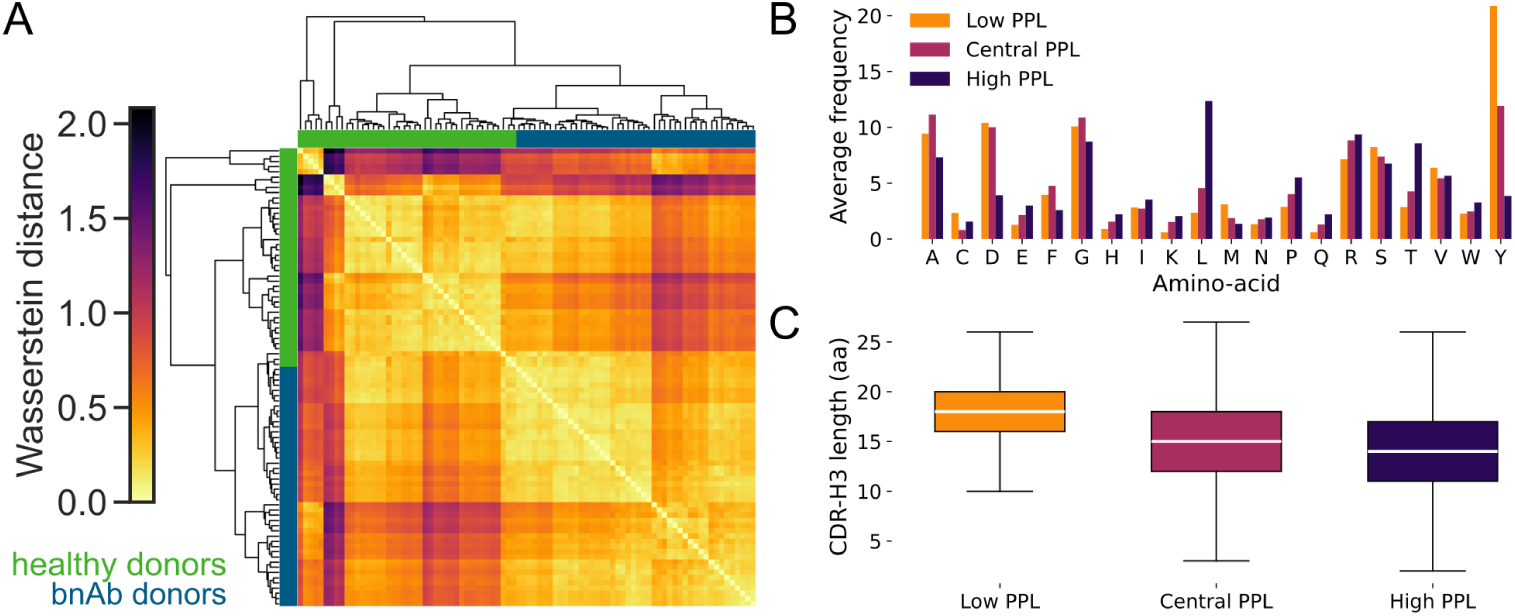
Pseudo-perplexity landscape of antibody repertoires quantified with H3BERTa. (A) Hierarchical clustering of donor repertoires based on pair-wise Wasserstein distances between their H3BERTa pseudo-perplexity distributions. Columns and rows correspond to individual repertoires; colour intensity reflects the distance scale shown at left (yellow = similar, purple/black = dissimilar). Side annotation: green, healthy donors; blue, donors who elicited broadly neutralizing antibodies. (B,C) Sequence features of CDR-H3s partitioned into low-, central- and high-perplexity (PPL) thirds (orange, magenta and dark violet, respectively). (B): mean amino-acid frequency in each bin. (C) Box-plots of CDR-H3 length for Low, central and high PPL (∼14 aa) sequences.

Within the healthy group, five distinct clusters were observed, each with low internal variability, indicating a higher degree of repertoire similarity within clusters (Figure 3A and Figure S3B). The largest cluster included approximately half of the healthy donors. Two additional clusters, each comprising approximately one-quarter of the healthy individuals, showed modest divergence from the main group. Finally, two small clusters, each containing only a few donors, stood out with distinctly different pseudo-perplexity profiles from the rest of the healthy cohort and from all bnAb donors. These outlier groups may reflect unusual immune histories or idiosyncratic repertoire development.

In the bnAb cohort, most donors grouped into two primary clusters with internally consistent pseudo-perplexity profiles. The intra-cluster distances were similar to those in the healthy group, whereas the average distance between the two bnAb clusters was higher, thus contributing to an overall increase in within-group variability (Figure S3). Interestingly, three healthy donors co-clustered with bnAb repertoires (Figure 3A), indicating a partial convergence or overlap in sequence plausibility profiles. These cases may reflect atypical repertoire features within the healthy population, potentially due to uncharacterized immune stimulation that share aspects of the bnAb response.

### 2.0.2 Differences in the distributional tails of pseudo-perplexity

Building on the observed clustering of repertoire-level pseudo-perplexity profiles, we next investigated whether these differences could be localized to specific regions of the distribution. Visual inspection of the merged PPL distributions revealed broad overlap between healthy and bnAb repertoires, particularly in the central range (Figure S4). The 5th–95th percentile intervals were nearly identical between groups (Healthy: [2.22, 16.61]; bnAb: [2.23, 16.41]), suggesting that any group-specific divergence may lie in the distributional tails. To test this hypothesis, we focused on the lower and upper tails of the PPL distributions, defined as sequences falling outside the central 90% range. First, the non-parametric Kolmogorov–Smirnov (KS) test was applied separately to the upper and lower tails. Both comparisons revealed statistically significant deviations from the null hypothesis (Upper tail: stat = 0.032, *p <* 0.001; Lower tail: stat = 0.057, *p <* 0.001), indicating that the shape of the distribution in the extremes differs between groups, with the effect more pronounced in the lower tail. Importantly, directional testing using the greater and less alternatives showed that, in the upper tail, healthy repertoires tend to exhibit higher PPL values than diseased repertoires. To complement this, the Mann–Whitney U test was used to assess whether the tails differed in location. This test also revealed significant group-level differences (Upper tail: *U* = 2.41×^10^, *p <* 0.001; Lower tail: *U* = 2.18 × 10^10^, *p <* 0.001), confirming that the group differences are concentrated in the tails rather than the center of the distribution. Together, these results demonstrate that the observed differences are concentrated in the tails of the distribution and are directionally biased, with healthy repertoires showing higher PPL in the upper tail while the lower tail differences reflect an opposite or more complex pattern.

### 2.0.3 Biochemical characterization of perplexity-defined CDR-H3 regions

To gain biochemical insights into the sequence properties associated with the perplexity groups, we analyzed the amino acid composition and CDR-H3 loop length within the Low, Central, and High PPL groups (lowest 5%, middle 90%, and highest 5% of perplexity values, respectively). Amino acid composition varied systematically across these PPL groups (Figure 3B). Loops categorized as Low PPL were significantly enriched in tyrosine (Y), an aromatic residue frequently involved in antigen binding. In contrast, High PPL loops showed a marked increase in the hydrophobic amino acid leucine (L), Threonine (T), along with more moderate enrichments in the conformationally restricted proline (P). These distinct changes in composition indicate that perplexity values reflect underlying biochemical profiles of CDR-H3 sequences.

A clear trend in CDR-H3 loop length was also observed across the perplexity-defined groups (Figure 3C, Figure S5). Low-PPL sequences were typically the longest (median = 17–18 residues), followed by Central-PPL (median = 15 residues), and High-PPL sequences, which were the shortest (median = 14 residues). Notably, H3BERTa consistently assigned central or high PPL scores to rare ultra-long CDR-H3 sequences. For example, 322 out of 8,649,880 total sequences exceeded 50 amino acids in length, with the longest reaching 77 residues, confirming their atypicality from the model’s learned distribution.

To evaluate whether model-derived perplexity correlates with broader biochemical properties, we analyzed net charge at physiological pH (7.0) and hydrophobicity using the GRAVY (Grand Average of Hydropathy) score for sequences within the Low-, Central-, and High-PPL bins (Figure S6 and Table 1). Net charge differed significantly across all groups. Low-PPL sequences were significantly less charged than both Central, whereas High-PPL sequences carried more charge than Central ones. While modest, the hydrophobicity differences were also statistically significant across all pairwise comparisons.

**Table 1.** Mann–Whitney U test results for Net Charge at pH 7 and Hydrophobicity (GRAVY) across PPL groups.

| Comparison | Property | U statistic | p-value |
| --- | --- | --- | --- |
| Low PPL vs Central PPL | Net Charge @ pH 7 | $1.26 \times 10^{12}$ | $p < 0.001$ |
| | Hydrophobicity (GRAVY) | $1.69 \times 10^{12}$ | $p < 0.001$ |
| Low PPL vs High PPL | Net Charge @ pH 7 | $4.71 \times 10^{10}$ | $p < 0.001$ |
| | Hydrophobicity (GRAVY) | $5.09 \times 10^{10}$ | $p < 0.001$ |
| Central PPL vs High PPL | Net Charge @ pH 7 | $1.16 \times 10^{12}$ | $p < 0.001$ |
| | Hydrophobicity (GRAVY) | $8.12 \times 10^{11}$ | $p < 0.001$ |

### 2.1 bnAbs classifier

Building on our repertoire-level analyses, we next investigated whether H3BERTa could also capture functional differences at the single-sequence level. Specifically, we evaluated whether its learned representations could support the classification of individual CDR-H3 loops based on their HIV-1 neutralization capacity. To this end, a labeled dataset of 390 experimentally validated CDR-H3 sequences, with an equal number of broadly neutralizing and non-neutralizing sequences was constructed. The bnAb sequences were sourced from the National Laboratory’s CATNAP database [32], a curated repository of HIV antibody neutralization data. The nnAb class combined CDR-H3s from HIV-infected individuals without broadly neutralizing activity and carefully selected “hard negatives” from healthy donors in the OAS database. These hard negatives, which share high sequence similarity with known bnAbs, were included to create a more challenging and realistic classification task. Despite this, the bnAb group exhibited a broader and right-shifted CDR-H3 length distribution relative to nnAbs, consistent with the known enrichment of long loops in bnAbs (Figure S7).

#### 2.1.1 Support Vector Machine

We trained a linear-kernel Support Vector Machine (SVM) using H3BERTa-derived embeddings as input features. This simple and interpretable model served as an initial probe to test whether the learned representations encode sufficient discriminative information for distinguishing bnAbs from nnAbs at the sequence level.

Model performance was first evaluated on a validation set, achieving an overall accuracy of 92% on the bnAbs class and a macro-averaged F1-score of 0.92 (Table 2), indicating balanced performance across classes.

**Table 2.** SVM performance on validation and test sets. Metrics are reported overall and per class. Acc. = Accuracy, P = Precision, R = Recall, F1 = F1-score.

| Set | Acc. | bnAbs |  |  | nnAbs |  |  |
| --- | --- | --- | --- | --- | --- | --- | --- |
|  |  | P | R | F1 | P | R | F1 |
| Validation | 92.0% | 1.00 | 0.83 | 0.91 | 0.86 | 1.00 | 0.92 |
| Test | 93.0% | 0.95 | 0.90 | 0.92 | 0.90 | 0.95 | 0.93 |

On an independent test set, performance remained consistent, with an overall accuracy of 93% and a macro F1-score of 0.92 (Table 2). This corresponded to correct classification of 95.0% of nnAbs and 90.0% of bnAbs, with a slightly higher false negative rate observed for bnAbs (10.0%) compared to nnAbs (5.0%). Visualization of the decision boundary in Principle Component Analysis (PCA)-reduced H3BERTa embedding space confirmed that most sequences were linearly separable, with misclassifications occurring primarily near the margin (Figure 4A).

**Fig. 4.**
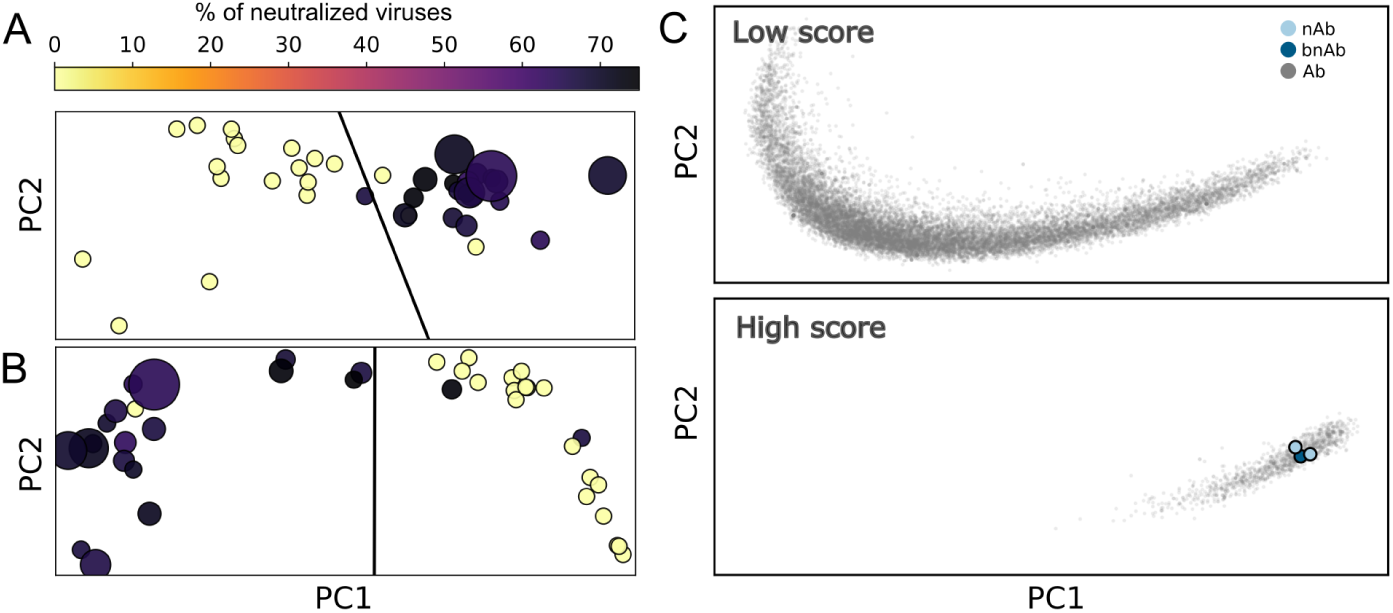
Antibody neutralization performance visualized through embedding-based decision boundaries and repertoire projection. (A, B) Decision boundaries of the SVM classifier trained on embeddings from (A) H3BERTa and (B) GAN-H3BERTa. Each dot represents an antibody of the test set. Dot size reflects the size of the viral panel on which the antibody was tested, while the color indicates the neutralization breadth, i.e., the number of viruses neutralized by the antibody divided by the total number of viruses in the assay. (C) PCA projection of GAN-BERT embeddings from an antibody repertoire of a donor with neutralizing serum [33]. Each dot represents an antibody from the donor’s repertoire. The top panel shows antibodies assigned low neutralization scores by the GAN-H3BERTa model, while the bottom panel shows those assigned high scores. Blue and light blue dots correspond to antibodies predicted and experimentally characeterized bnAbs and nAbs [33], respectively, and were also correctly identified as such by the GAN-H3BERTa approach.

#### 2.1.2 GAN-H3BERTa

To explore whether a deep learning approach could reduce the false positive rate observed with the SVM, we adopted a semi-supervised framework suited to settings with limited labeled data. Specifically, we employed GAN-BERT [34], which integrates a pre-trained language model with adversarial fine-tuning to exploit both labeled and unlabeled data during training. Unlike conventional supervised models, GAN-BERT is designed to learn from a small number of labeled examples while also incorporating distributional information from a larger pool of unlabeled sequences. In our adaptation, the original BERT encoder was replaced with H3BERTa, and the model was fine-tuned using labeled CDR-H3 sequences and unlabeled sequences from the repertoires of HIV-infected individuals (Figure 1B). To construct the unlabeled set, we sampled CDR-H3 loops from antibody repertoires of HIV-positive donors with varying degrees of neutralization breadth. To enrich for functionally mature antibodies, only heavy-chain sequences with *<* 85% V gene identity were retained, consistent with the high levels of somatic hypermutation typically observed in HIV-related immune responses.

The GAN-H3BERTa model achieved 83.3% accuracy on the validation set (Matthews correlation coefficient (MCC) 0.67, Area Under the Curve (AUC) 0.83), showing good class balance (Table 3). On the independent test set, accuracy rose to 85.0% (MCC 0.71, AUC 0.85), with bnAbs detected at high sensitivity (recall 0.95) while maintaining acceptable precision (0.79)(Table 3). Overall, the model demonstrated strong generalization and a good balance between sensitivity and precision across classes.

**Table 3.** GAN-H3BERTa performance on validation and test sets. Metrics are reported overall and per class. Acc. = Accuracy, MCC = Matthews correlation coefficient, AUC = Area under the ROC curve, P = Precision, R = Recall, F1 = F1-score.

| Set | Acc. | MCC | AUC | bnAbs |  |  | nnAbs |  |  |
| --- | --- | --- | --- | --- | --- | --- | --- | --- | --- |
|  |  |  |  | P | R | F1 | P | R | F1 |
| Validation | 83.3% | 0.67 | 0.83 | 0.80 | 0.89 | 0.84 | 0.88 | 0.78 | 0.82 |
| Test | 85.0% | 0.71 | 0.85 | 0.79 | 0.95 | 0.86 | 0.94 | 0.75 | 0.83 |

To evaluate how adversarial fine-tuning reshaped the embedding space, we trained a linear SVM on the CDR-H3 embeddings extracted from the fine-tuned H3BERTa encoder within GAN-H3BERTa. This allowed comparing the separability of antibody classes in the fine-tuned space to that of the original, static H3BERTa embeddings. On the validation set, the SVM trained on GAN-BERT embeddings achieved an accuracy of 92% and a macro-averaged F1-score of 0.92, indicating strong and balanced performance across classes. Misclassifications were limited, with less than 6% of bnAbs labeled as nnAbs, and 11% false positives in the nnAb group. These results were consistent on the test set, where accuracy reached 93% and the macro F1-score remained at 0.92. Class-level precision and recall also remained balanced, with slightly improved recall for nnAbs and precision for bnAbs. Overall, both the validation and test sets confirmed that adversarial fine-tuning led to embeddings that support clearer class separation. Although both SVMs showed structured decision boundaries (Figure 4A–B), the latent space learned by GAN-H3BERTa appeared to be more compact and further from the decision margin, suggesting that semisupervised training helped shape a more robust and discriminative feature space.

Finally, to assess model performance under more realistic conditions, we applied both the SVM and GAN-H3BERTa classifiers to the complete B cell repertoire from an HIV-1–infected donor known to produce broadly neutralizing antibodies [33], which includes experimentally characterized antibodies. This analysis allows evaluating whether the models can correctly identify known bnAbs while estimating the overall specificity in a full repertoire. Overall, the SVM predicted 31.1% sequences as bnAbs, while the GAN-H3BERTa approach reduced this slightly to 28.0% sequences, reflecting a modest improvement in specificity. Importantly, both models correctly classified the experimentally validated antibodies, one bnAb and two neutralizers, as bnAbs. In latent space, these sequences clustered within high-confidence bnAb regions (Figure 4C), and HIV-1–specific sequences more broadly localized to the same region, suggesting that H3BERTa captures relevant antigen-associated features even in complex, unfiltered repertoires.

## 3 Discussion

The identification of antigen-specific antibodies within the vast and heterogeneous B cell receptor repertoire remains a major challenge in immunology, vaccinology, and therapeutic antibody discovery. Although next-generation sequencing has enabled unprecedented access to antibody repertoires, extracting functionally relevant candidates from this immense sequence space remains difficult, particularly in the absence of antigen-specific labeling. Recent computational approaches, including machine learning–based models, have sought to identify patterns in antibody sequences[2, 18–20]; most rely on full-length variable-region sequences. Self-supervised pretraining of such models can capture broad sequence representations, However, the larger parameter space and broader sequence coverage increase the amount of labeled data and computational resources required for downstream fine-tuning, since the model must learn to focus on task-relevant features within a much richer representation space.

We hypothesized that focusing exclusively on the hypervariable CDR-H3 region could retain much of the relevant immunological signal while reducing the complexity of the representation space. This would enable effective downstream analysis even in low-label or fragmentary data settings. We developed H3BERTa, a transformer-based language model trained on over 18 million CDR-H3 sequences, to test whether this hypervariable region alone could capture meaningful immunological signals in antibody repertoires. Our findings show that despite the reduced input scope, H3BERTa successfully recapitulated many biologically coherent repertoire-level features. The most prominent structure in H3BERTa’s latent space was clustering by J-gene usage, reflecting the contribution of IGHJ segments to the C-terminal residues of CDR-H3 and, more broadly, the influence of J-gene choice on loop length and amino acid composition. These sequence-level effects shape the physicochemical properties of the CDR-H3 loop that are relevant for antigen recognition [35].

To assess whether H3BERTa embeddings capture functional signals beyond sequence composition, we tested whether the B cell maturation state could be inferred from the latent representations using a simple linear discriminant analysis. Although maturation is associated with the accumulation of somatic hypermutations, CDR-H3 sequences are short and highly diverse, making extraction of relevant information challenging. The ability to discriminate maturation states from these representations indicates that H3BERTa encodes functionally meaningful features within the hypervariable region, supporting its applicability for repertoire-level analyses. Although some separation between populations may arise from J-gene usage and other maturation-linked sequence features, the fact that these distinctions can be captured using only CDR-H3 sequences underscores the model’s capacity to represent biologically relevant repertoire information.

H3BERTa also enables fine-grained comparison of pseudo-perplexity distributions, providing a complementary view of sequence diversity and maturation within antibody repertoires. Although the overall pseudo-perplexity profiles of healthy and HIV-1–derived repertoires largely overlap, both statistical testing and distributional metrics revealed consistent differences, particularly at the extremes. It is known that next-generation sequencing captures only a small fraction of the total B-cell repertoire [36]. Nonetheless, because these distributional differences are observed systematically across samples, they are unlikely to arise from random sampling effects and may reflect underlying biological variation. Further work will be needed to determine the immunological processes contributing to these patterns. Low-perplexity sequences tend to have longer CDR-H3 loops enriched in tyrosine residues, features often associated with greater structural complexity and affinity maturation [37]. In contrast, high-perplexity sequences more frequently display shorter, leucine-rich CDR-H3s, consistent with less mature or more flexible binding configurations [38]. The presence of these distributional differences between healthy and HIV-1 repertoires is consistent with altered B-cell selection and maturation under chronic antigen exposure [39–42].

Importantly, these divergences cannot be explained solely by CDR-H3 length or amino-acid composition, indicating that H3BERTa captures additional sequence-level features beyond simple compositional biases. We also observed that sequences with higher net positive charge tend to exhibit greater pseudo-perplexity, implying that electrostatic properties influence model uncertainty. This observation aligns with reports that affinity maturation and antigen engagement involve changes in surface charge and loop dynamics [43]. Together, these results suggest that H3BERTa’s pseudo-perplexity distributions encode biologically relevant biophysical and evolutionary information, offering a sensitive statistical framework for comparing antibody repertoires even when only limited sequence data are available.

The effectiveness of H3BERTa embeddings in downstream classification tasks demonstrates that the model captures discriminative, task-relevant information within the CDR-H3 region. Even a simple linear SVM achieved strong separation between neutralizing and non-neutralizing antibodies, highlighting the biological richness of the learned representations. When the SVM classifier was applied to full B-cell repertoires, its limitations became evident. Although it performed well on balanced validation sets, it generated many false positives in realistic repertoires, where broadly neutralizing antibodies (bnAbs) constitute only a small fraction of all sequences [14]. This outcome illustrates a common challenge in repertoire-level prediction: extreme class imbalance and biological heterogeneity can cause apparent model performance to degrade when moving from curated datasets to natural repertoires. To mitigate this, we explored a semi-supervised GAN-BERT framework that fine-tunes H3BERTa’s embedding space using limited labeled data. This approach improved the separability of bnAb sequences and modestly reduced false positives, demonstrating the potential of semi-supervised strategies for antibody classification under data-scarce conditions. While additional data and further methodological development will be required to achieve stronger discrimination, these results suggest that integrating generative and discriminative learning could be a fruitful direction for future repertoire modeling efforts.

In sum, these results demonstrate that H3BERTa captures not only germline- and structure-driven aspects of antibody repertoires but also subtler evolutionary and biophysical signals associated with affinity maturation and immune history. By distilling immunologically meaningful representations from the hypervariable CDR-H3 region, H3BERTa provides a compact yet informative framework for analyzing repertoire dynamics across healthy and diseased donors. Such representations hold promise for advancing antibody discovery, vaccine design, and longitudinal immune monitoring. Future work will focus on evaluating H3BERTa across broader datasets and disease contexts, assessing its capacity to predict antigen specificity and neutralization breadth directly from sequence, and refining the GAN-BERT framework to improve detection of rare, therapeutically relevant antibodies in large-scale repertoires.

## 4 Materials and Methods

### 4.1 H3BERTa

H3BERTa is a transformer-based language model designed specifically for CDR-H3 loop sequences of B-cell receptors. The model was trained on sequences sourced from the Observed Antibody Space (OAS) database [24].

#### Dataset

The H3BERTa training dataset was constructed by extracting the CDR-H3 sequences from the OAS database [24], specifically selecting entries from healthy human donors with no documented history of vaccination or disease, and including all available B cell sources (Figure 1A). IgG and IgA isotype sequences were extracted and CDR-H3 loops were identified using IMGT annotations. To reduce redundancy, the sequences were clustered at 70% identity using MMseqs2 [44], retaining only the cluster centroids. Sequences containing more than two ambiguous amino acids (e.g., ’X’) or with CDR-H3 loops shorter than three residues were excluded, resulting in a final dataset of approximately 18 million sequences. The remaining centroids were further clustered at 30% identity to create three distinct splits: training (80%), validation (10%), and testing (10%). To ensure data consistency, all sequences within a 30% identity cluster were assigned to the same split, with larger clusters preferentially allocated to the training set.

#### Model

We used the RoBERTa architecture [23] as the base model and conducted an extensive architecture search to explore variations in model depth and complexity. The search involved adjusting the hidden size (from 512 to 768), intermediate size (from 2048 to 3072), the number of attention heads (from 4 to 12), and the number of attention layers (from 4 to 12). This exploration resulted in a range of model sizes, with the number of trainable parameters varying from 16 million to 86 million. The final model configuration features a hidden size of 768, an intermediate size of 3072, 12 attention heads, and 12 attention layers, totaling 86 million parameters. Sequences were tokenized at the amino acid level, with a maximum sequence length fixed at 100 residues.

#### Training

The batch size was set to 1024, and we tested learning rates of 1*e* − 3, 5*e* − 5, and 1*e* − 6, with 5*e* − 5 emerging as optimal. Training was performed using the AdamW optimizer with a weight decay of 0.1, and took approximately one month on a single A100 80G NVIDIA GPU to reach convergence at epoch 113.

#### Blosum score

During the masking objective for each masked position in the sequence, we compared the model’s top prediction to the original amino acid and retrieved the corresponding substitution score from the BLOSUM62 matrix (via Biopython’s Bio.Align.substitution matrices module). If a given pair of amino acids was not present in the matrix, for example, due to the presence of non-canonical residues or special tokens like [UNK], a penalty score of -5 was assigned. The BLOSUM scores were aggregated across all masked positions to obtain an overall measure of prediction quality that accounts for biochemical similarity, rather than exact matches alone.

### 4.2 Classifier

#### bnAbs dataset

We compiled our labeled dataset using information from multiple studies available in the Los Alamos National Laboratory’s HIV Database (CATNAP) [32]. Our selection was restricted to human-derived data.

We categorized antibodies into two groups:

- Broadly neutralizing antibodies (bnAbs): Antibodies with an average neutralization detection rate exceeding 25%.
- Non-neutralizing antibodies (nnAbs): Antibodies with no detectable neutralization activity.

We retrieved available nucleotide sequences and processed them using the Python package PyIR [45], isolating the CDR-H3 based on IMGT annotations. To mitigate initial class imbalance and introduce challenging negative examples, we enriched the nnAbs category by including CDR-H3 sequences from healthy individuals from OAS that shared over 70% sequence identity with bnAbs retrieved using blastp with the following default parameters -evalue 2e-5 -outfmt 6 -word size 2 -threshold 11 -matrix PAM30 -window size 40 -comp based stats 0. This resulted in a perfectly balanced dataset comprising 195 sequences per class. To prevent information leakage, we used MMseqs2, specifically the Linclust algorithm with default parameters, to cluster sequences within each antibody category prior to splitting. The resulting clusters were then used to partition the dataset into training (80%), validation (10%), and test (10%) subsets using a standard split strategy. All sequences within a given cluster were assigned to the same dataset split, with larger clusters preferentially allocated to the training set to ensure sufficient representation, while preserving the independence of the validation and test sets. Additionally, each split was randomly shuffled to eliminate potential ordering effects before training and evaluation.

#### Support Vector Machine

We employed the pretrained H3BERTa model, which was trained on healthy antibody repertoires, as a fixed embeddings extractor. Each input CDR-H3 sequence was tokenized at the single-residue level and fed through the model. To obtain a fixed-length embedding for each sequence, we applied mean pooling over the final hidden states, averaging only over positions corresponding to valid (non-padded) tokens as indicated by the attention mask. The resulting embeddings, representing contextualized sequence-level features, were subsequently used to train a linear Support Vector Machine (SVM) classifier. The classifier was implemented using the scikit-learn library, trained on the designated training set, and evaluated on a held-out test set to assess generalization performance. For comparison, we repeated the entire procedure using the pretrained AntiBERTa model as a fixed embedding extractor.

#### GAN-BERT

For the detection of bnAbs, we employed the GAN-BERT approach [34], an adversarial classifier that combines a pre-trained language model with adversarial training. This method leverages both a small labeled dataset and a larger unlabeled dataset.

#### Labeled Dataset

For the labeled dataset, we used the annotated data, that was employed to train, validate and test the SVM classifier.

#### Unlabeled Dataset

Antibody heavy-chain repertoire sequences were obtained from a previously published dataset [29], comprising 139 individuals with chronic HIV-1 infection (46 with broad neutralization breadth, 50 with limited breadth) and 43 HIV-uninfected controls. Repertoires were labeled according to donor-level neutralization phenotype: bnAbs (broad neutralization), nAbs (limited neutralization), and nnAbs (HIV-negative controls/healthy donors). In this study, we employed only bnAbs and nnAbs phenotypes.

Sequences were filtered by V gene germline identity, retaining only those below 85%. Unique CDR-H3 loops were extracted and used as a dataset.

Two configurations of the unlabeled dataset were constructed. In the balanced version, approximately 10,000 sequences from each of the bnAb and nAb groups were combined with ∼20,000 randomly sampled sequences from the nnAb group. In the HIV-positive-only version, only bnAb and nAb repertoires were included (∼10,000 sequences per group), assuming the majority of sequences in these repertoires to be non-neutralizing.

#### Model

We implemented a semi-supervised learning model based on the GAN-BERT framework [34], utilizing the Hugging Face Transformers library. Our model employs H3BERTa as the language model backbone and integrates a Generative Adversarial Network (GAN) consisting of a generator and a discriminator.

The generator (G) is a multilayer feedforward neural network. It takes as input a random vector of size 1000, sampled from a normal distribution, and produces ”fake” embedding vectors of dimension 768, matching the dimensionality of the real H3BERTa embeddings. The generator architecture consists of 1 hidden fully connected layer with 768 units, followed by a LeakyReLU activation function (with negative slope 0.01) and a dropout layer with rate 0.1.

The discriminator (D) is a multilayer feedforward neural network that takes as input either a real embedding from H3BERTa (corresponding to a CDR-H3 sequence) or a fake embedding generated by G. Its output is a three-way classification, distinguishing between the nnAbs and bnAbs categories, as well as a third category representing the fake embeddings generated by G. The discriminator architecture consists of 2 fully connected layers with 768 units each, each followed by a LeakyReLU activation and dropout (rate 0.1).

The H3BERTa model was fine-tuned during training. The real embeddings used as input to the discriminator were obtained by passing the CDR-H3 sequences through the H3BERTa model. Both generator and discriminator were optimized using the Adam optimizer with a learning rate of 5e-5 and epsilon=1e-8. The model was trained across 1 GPU without a learning rate scheduler with a warmup proportion of 0.1.

#### Training

We conducted a hyperparameter search to optimize our model’s performance. The following hyperparameters were explored:

- Learning rates: 5 × 10*^−^*^5^ and 5 × 10*^−^*^6^.
- Hidden layers in both the generator and discriminator: 1, 2, and 3 layers.
- Generator input dimensionality (Noise size): 10, 50, 100, 500, and 1000.
- Dropout rate: 0.0, 0.1, 0.2, and 0.3 applied at the output layer of the discriminator.
- Batch size: 16, 32, 64, 128, and 256.

The final model was trained on a shuffled version of the balanced, unlabeled dataset described for the unsupervised objective, and on the labeled dataset introduced for the supervised objective. Sequences were tokenized at the amino acid level, with a maximum sequence length of 100 residues.

Training was conducted using adversarial objectives for both generator and discriminator, combined with a supervised cross-entropy loss applied to labeled data. The discriminator loss was defined as the sum of the supervised and unsupervised terms, while the generator loss included both adversarial and feature-matching components.

The optimal configuration, selected based on the lowest validation loss, comprised a batch size of 16, learning rates of 5*e* − 5 for both the generator and the discriminator, one hidden layer in the generator, two hidden layers in the discriminator, a noise vector size of 1000, and a dropout rate of 0.1. Model training was performed on a single NVIDIA A100 80GB GPU.

### 4.3 Evaluation

#### Single repertoire analysis

From the same unlabaled dataset used to train the classifier (Material and Methods 4.2), we selected one repertoire from each phenotype: one from the healthy group (id:700011206, number of not unique sequences: 113450) and one from the bnAb group (id: 702010293, number of not unique sequences: 115719).

#### Naive and Memory B-cells

The dataset used to extract embeddings of naive and memory B cells, consists of paired light and heavy chain antibody sequences extracted from the OAS database. We first downloaded all available human sequences. To ensure uniqueness, duplicates were removed based on the CDR3 region of the heavy chain. A balanced dataset was then created by randomly selecting 50,000 unique sequences from Memory B cells and another 50,000 from Naive B cells. A fixed random seed was applied to guarantee reproducibility.

#### Antibody therapeutics

All approved and phase 1–3 antibody therapeutics were retrieved from TheraSAbDab [28]. The origin of each therapeutic was determined based on its source infix and mapped to one of the following categories: Human (“u”), Humanized (“zu”), Chimeric/Humanized (“xizu”), Chimeric (“xi”), and Mouse (“o”). For the purposes of this analysis, therapeutics categorized as Chimeric/Humanized were excluded. A final dataset of 199 antibody therapeutics has been used.

#### Immune repertoires analysis

We employed again the same collections of repertoires from the single repertoire analysis (4.2), focusing specifically on uninfected individuals (healthy donors) and HIV-1-infected individuals with confirmed broadly neutralizing antibody (bnAb) responses. The dataset consisted of CDR-H3 immune repertoire sequences from 42 healthy and 46 HIV-1-infected donors. Unlike the classification task, no filtering or preprocessing steps were applied to the sequences prior to analysis. Pseudo-perplexity scores for all CDR-H3 sequences were computed using our H3BERTa model, and the resulting scores were subsequently analyzed to assess repertoire-level differences between the two groups.

#### PPL

To estimate the pseudo-perplexity of individual sequences using a masked language model (MLM) like H3BERTa, we followed the token-by-token masking approach introduced by [46]. Given a pre-trained MLM and a target sequence, each non-special token in the sequence was iteratively replaced with the model’s mask token. The masked sequence was then passed through the model to compute the log-probability of the true token at the masked position, conditioned on the rest of the sequence. This process was repeated for all tokens (excluding special tokens), and the final pseudo-perplexity was computed as the exponential of the negative average log-probability:

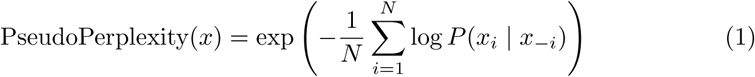

where *x_i_* is the true token at position *i*, and *x_−i_* denotes the sequence with the *i*th token masked. This method provides a token-level approximation of perplexity suitable for uni bidirectional models. The implementation was performed using PyTorch and HuggingFace Transformers. Gradients were disabled during inference.

## 5 Resource availability

All code, pretrained models, and curated datasets used in this study are openly available to support transparency, reproducibility, and community-driven development. Resources can be accessed via GitHub at the following repository: https://github.com/ibmm-unibe-ch/H3BERTa

## Acknowledgment

This work is supported by funds from the Helmut Horten Stiftung young researcher grant and the Swiss National Science Foundation (PCEFP3 194606)

## 7 Declaration of interests

The authors declare no competing interests.

### 7.1 Declaration of generative AI and AI-assisted technologies in the writing process

During the preparation of this manuscript, the authors employed generative AI tools, including ChatGPT (OpenAI) and Gemini (Google), for language refinement. These tools were used solely to improve clarity, coherence, and style. All AI-assisted content was carefully reviewed and substantively edited by the authors, who accept full responsibility for the accuracy and integrity of the final submitted work.

---

**Fig. S1.**
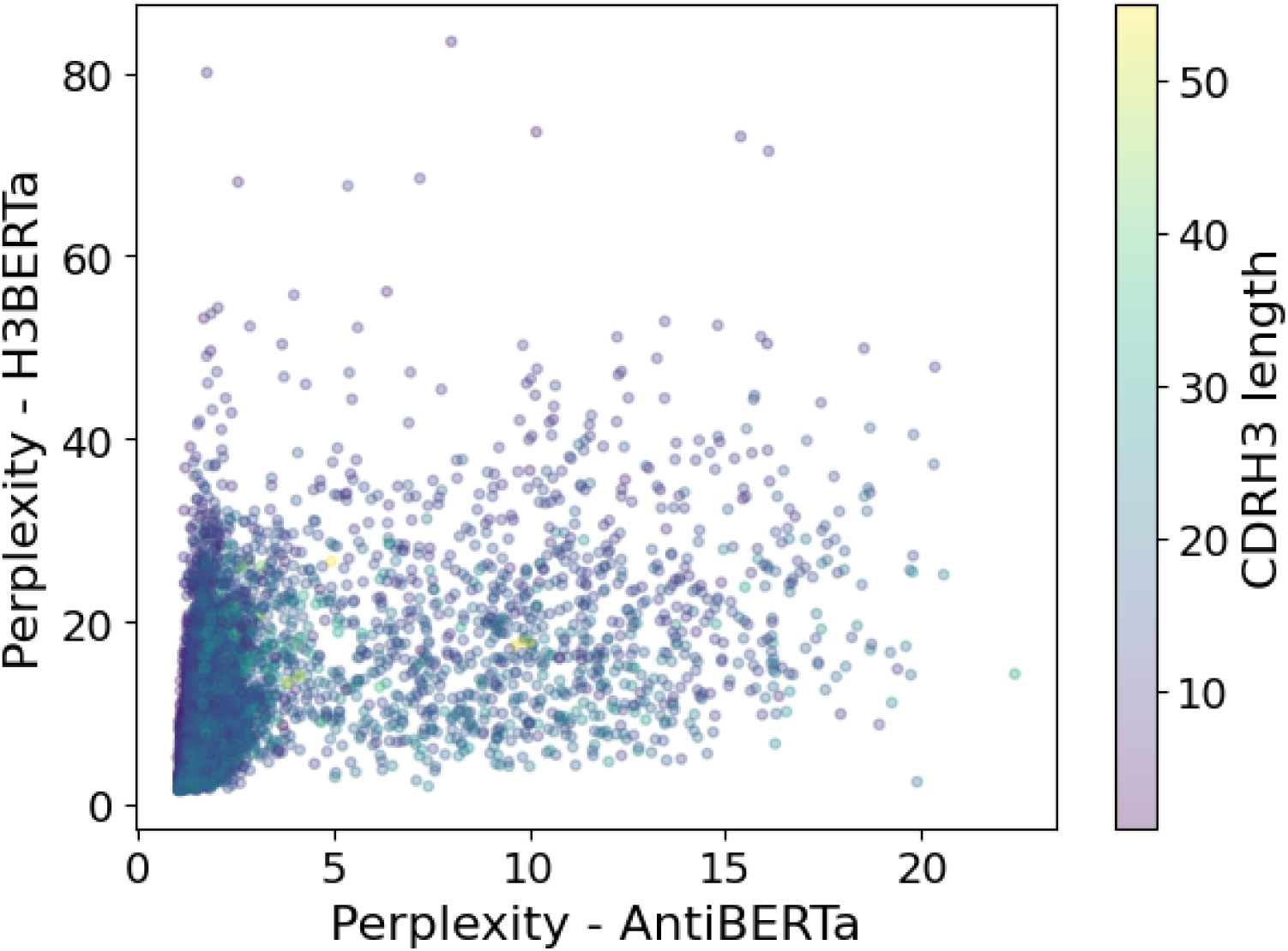
Comparison of perplexity values calculated with AntiBERTa (x-axis) and H3BERTa (yaxis) on a healthy repertoire, specifically on the heavy chain and CDR-H3 loop, respectively. Each point represents a unique CDR-H3 sequence: shifts to the right indicate an increase in perplexity for AntiBERTa, and shifts upwards indicate an increase for H3BERTa. Points are colored according to CDR-H3 length using a continuous colormap.

**Fig. S2.**
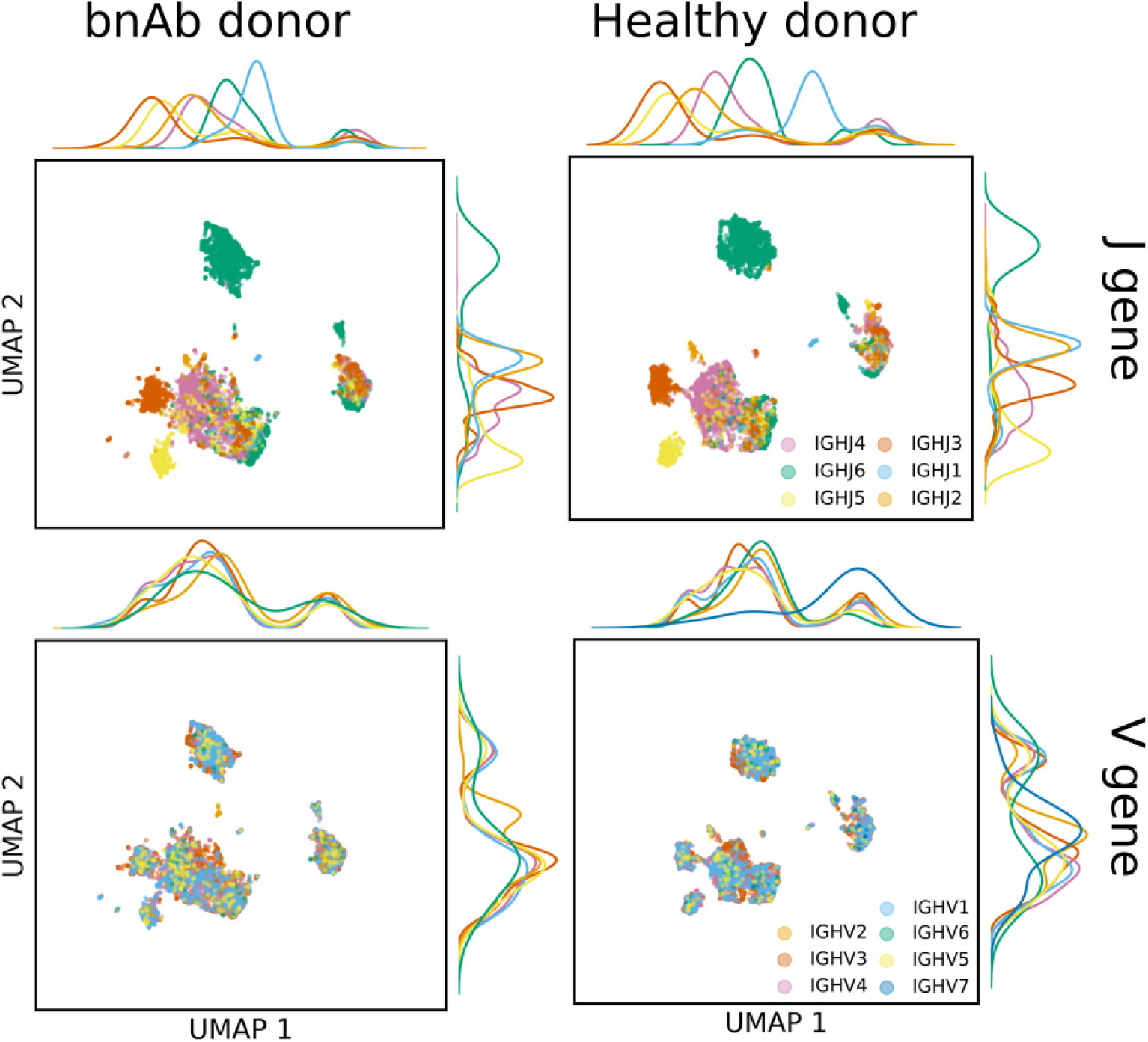
UMAP visualization of H3BERTa-derived embeddings for CDR-H3 loops from a bnAb donor (left) versus a healthy donor (right). In the top row, each point represents a single CDR-H3 embedding colored by IGHJ gene–segment usage; in the bottom row, embeddings are colored by IGHV gene–segment usage. Marginal ridge plots alongside each UMAP axis depict the density of individual Vand J-gene families.

**Fig. S3.**
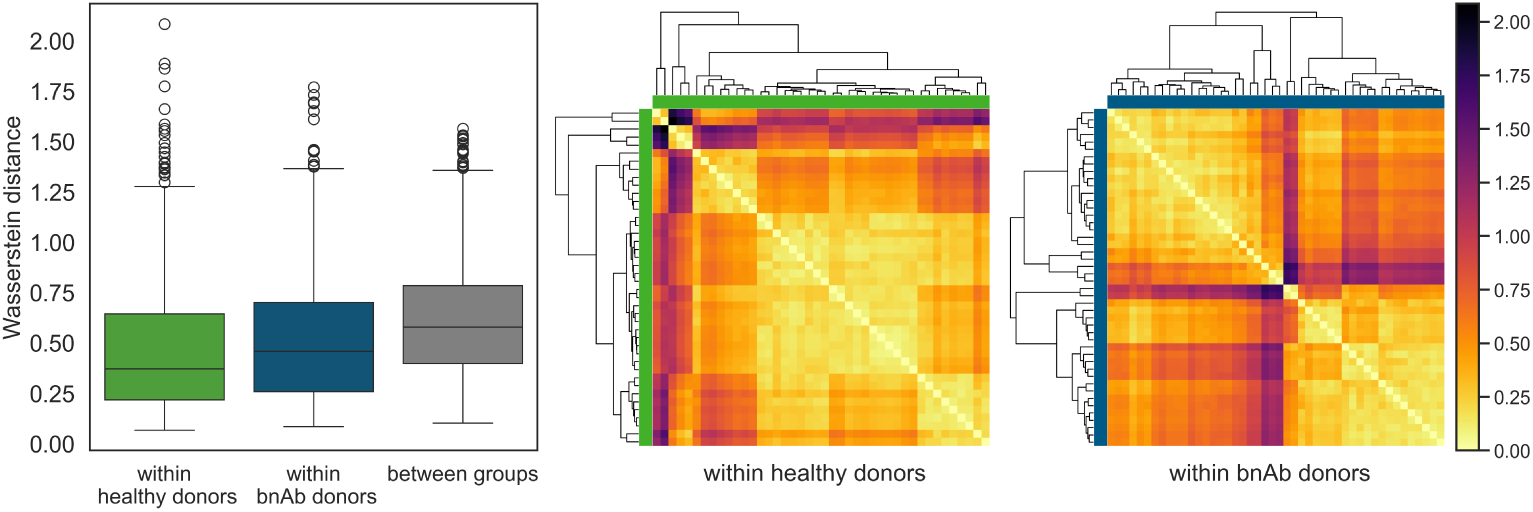
(A) Box plots of pairwise Wasserstein distances between CDR-H3 perplexity distributions from H3BERTA. Distances are shown for comparisons within healthy donors (green), within bNAb donors (blue), and between the two cohorts (grey). Lower values indicate more similar repertoires. Healthy donors display the lowest median distance, whereas inter-cohort comparisons are shifted upward, signalling a partial separation of the two perplexity landscapes. (B) Heatmaps of the same pairwise distances. The left panel includes only healthy donors; the right panel includes only bNAb donors. Samples are ordered by hierarchical clustering to highlight similarity patterns. Colours progress from yellow (0; high similarity) through orange to black (= 2; high dissimilarity). The healthy cohort forms a largely homogeneous block, while the bNAb cohort shows several darker off-diagonal patches, pointing to greater intra-group heterogeneity.

**Fig. S4.**
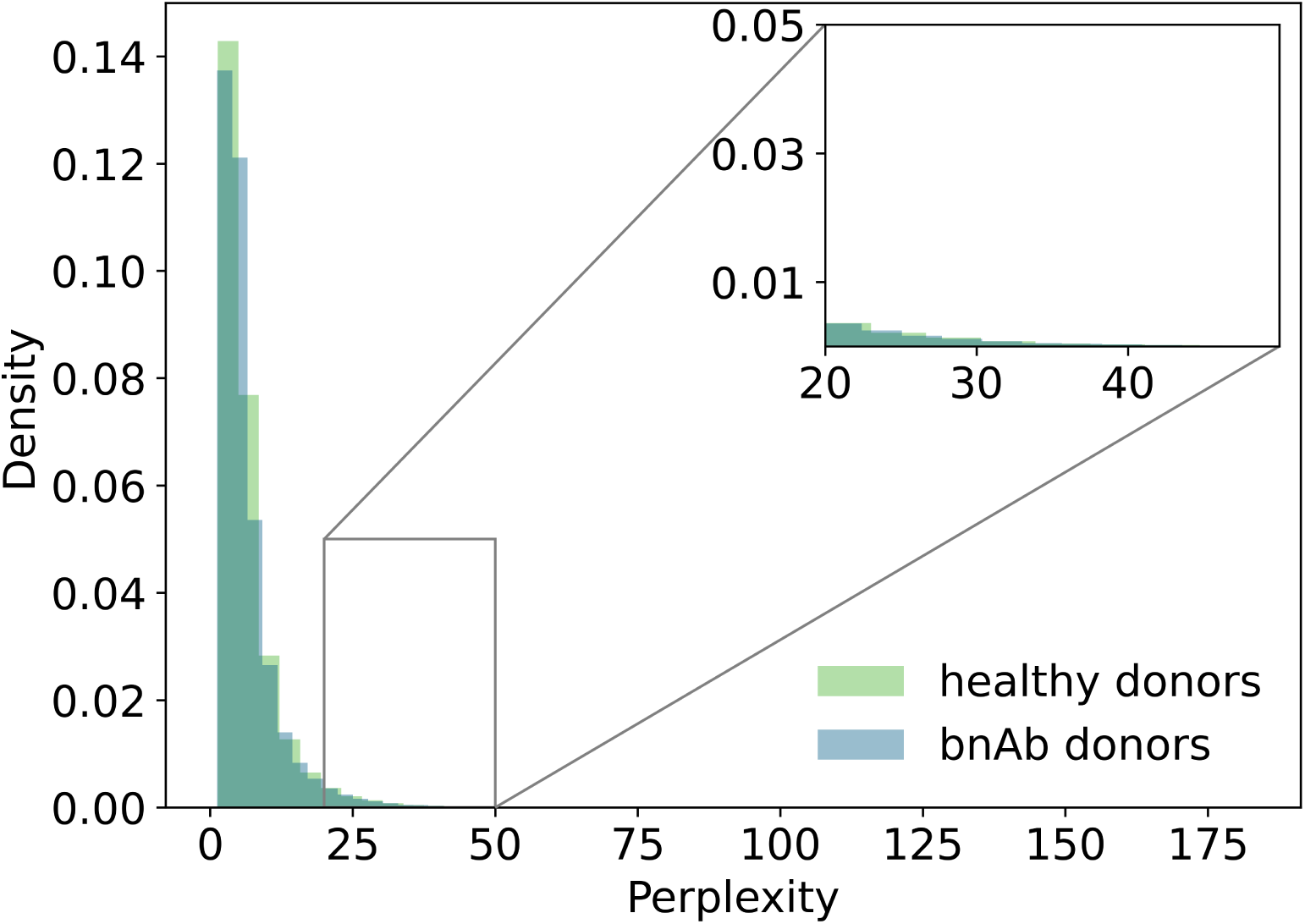
Perplexity distributions of CDR-H3 sequences from healthy and bnAb donors. Kernel density estimates show that both distributions are heavily right-skewed, with most sequences having low perplexity scores. An inset highlights the right tail of the distribution, where rare high-perplexity sequences are more visible.

**Fig. S5.**
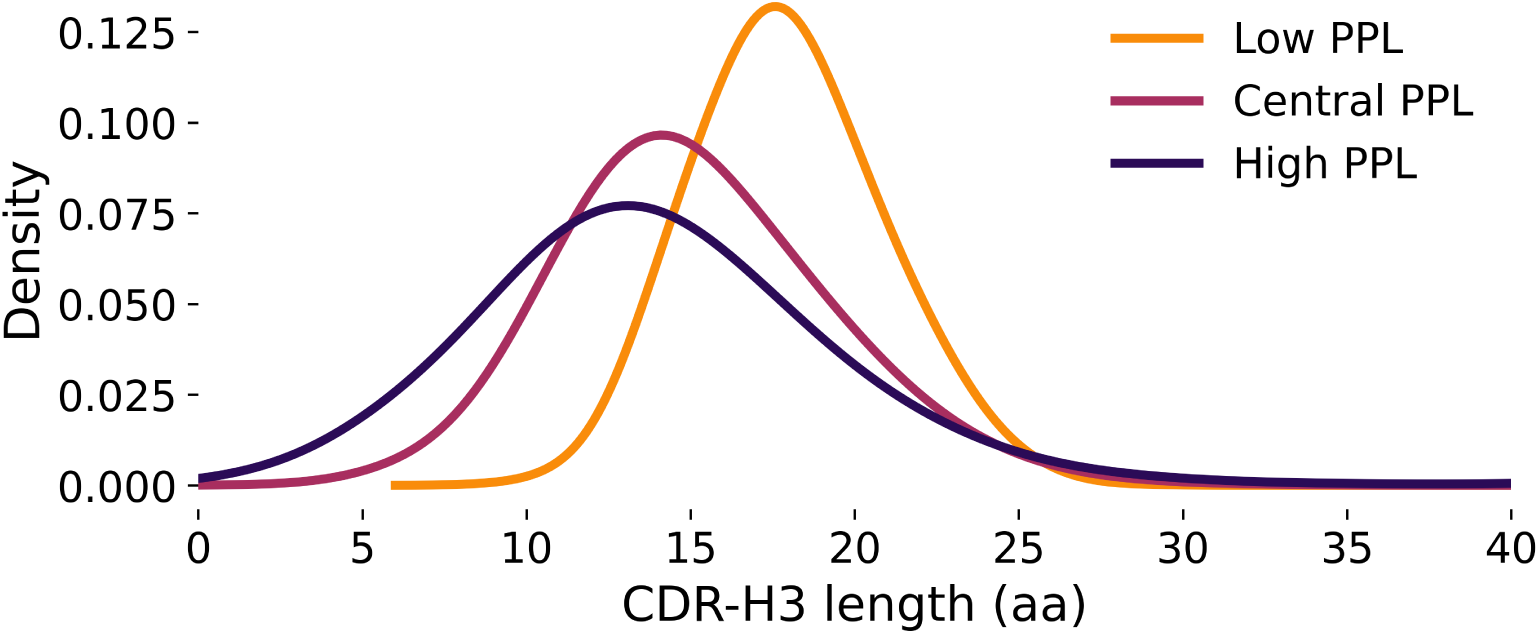
Length distribution of CDR-H3 sequences across perplexity-defined groups. Kernel density estimates of CDR-H3 amino acid lengths for sequences categorized into Low PPL (orange), Central PPL (magenta), and High PPL (dark purple) groups. While all groups display a peak in the 12–18 amino acid range, Low PPL sequences tend to be longer, indicating a potential association between lower model perplexity and naive CDR-H3 regions. CDR-H3 sequences with lengths greater than 40 amino acids were removed from the plot as outliers to improve the clarity of the density distribution.

**Fig. S6.**
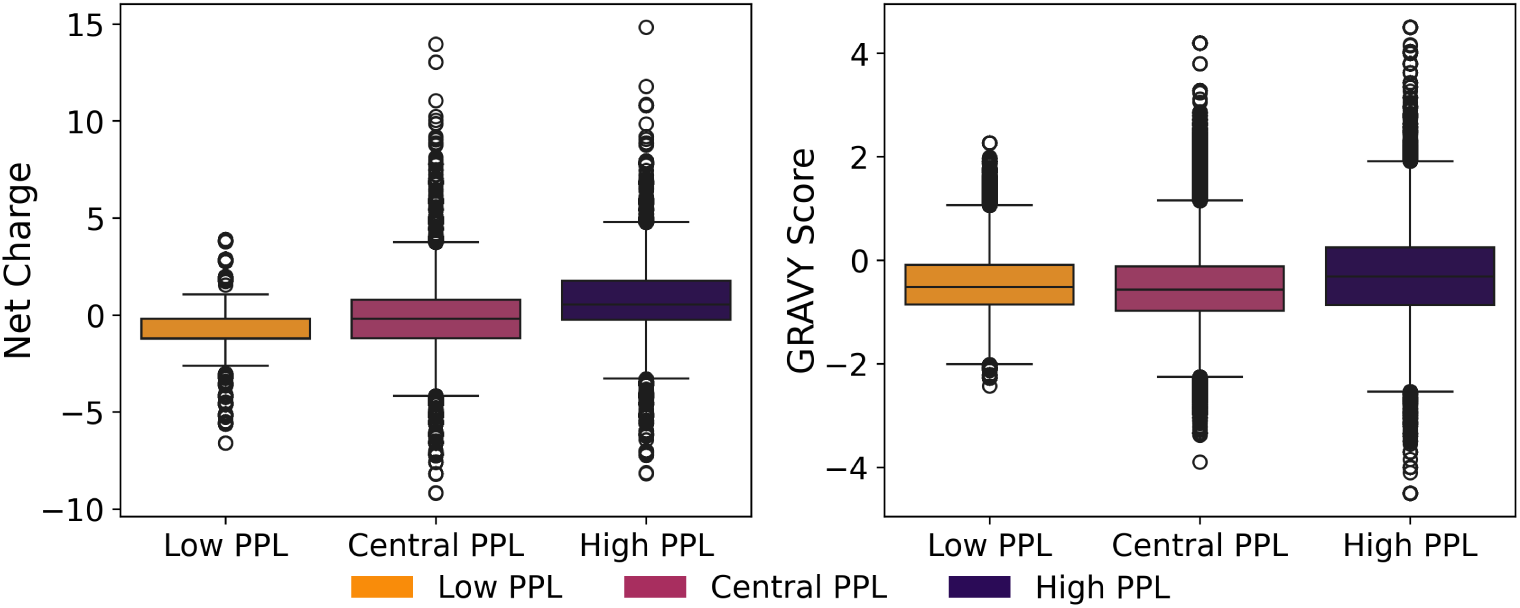
Biochemical properties of CDR-H3 sequences across perplexity-defined groups. Boxplots of (A) net charge at physiological pH (7.0) and (B) hydrophobicity (GRAVY score) for CDR-H3 sequences stratified by model-derived perplexity into Low, Central, and High PPL groups. Sequences in the Low PPL group exhibit significantly lower net charge compared to Central and High PPL groups (all *p <* 0.001, Mann–Whitney U test), suggesting a possible association between low perplexity and reduced electrostatic diversity. In contrast, GRAVY scores are relatively stable across groups, with only minor shifts despite statistically significant differences, indicating that overall hydrophobicity is not strongly differentiated by perplexity. Color-coded legends indicate PPL groups.

**Fig. S7.**
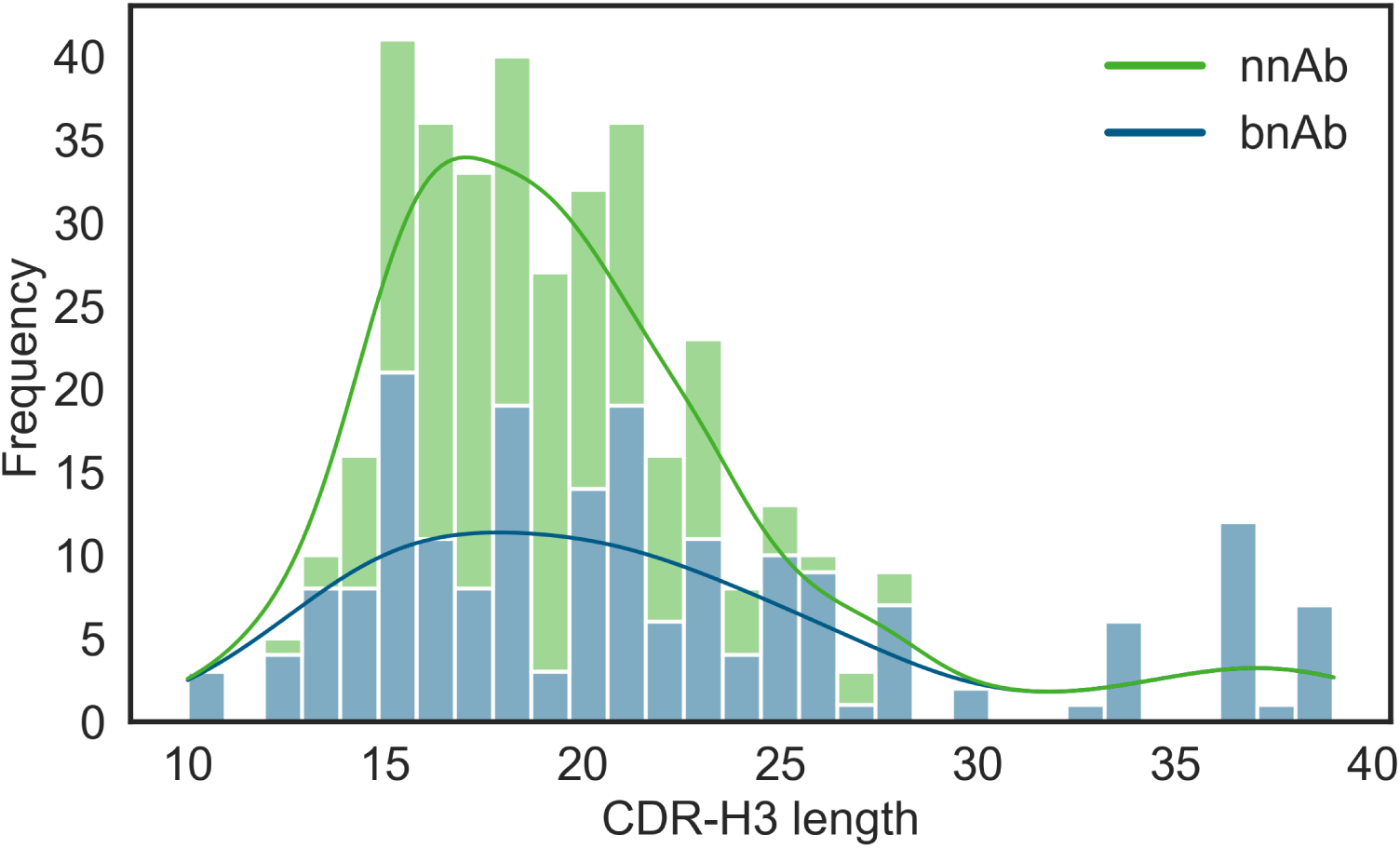
Length distribution of CDR-H3 loops in bnAb and nnAb repertoires. Histogram and kernel density estimates (KDE) of CDR-H3 amino acid lengths for broadly neutralizing antibodies (bnAbs, blue) and non-neutralizing antibodies (nnAbs, green). While both distributions peak in the 14–18 residue range, bnAbs tend to exhibit a broader and slightly right-shifted distribution, with a higher proportion of longer CDR-H3 loops compared to nnAbs.

